# GABA-Edited Magnetic Resonance Spectroscopy Deep Learning Quality Assessment Framework

**DOI:** 10.1101/2025.09.11.674310

**Authors:** Hanna Bugler, Roberto Souza, Ashley D. Harris

## Abstract

**Purpose:** Motivated by the need to improve GABA-edited magnetic resonance spectroscopy (MRS) quality, we developed a three-module framework to improve transient averaging based on quality. We hypothesized that training a deep learning (DL) to differentiate spectrum quality could improve transient averaging compared to traditional averaging.

**Methods:** The transient averaging framework was approached through three modules: (1) a continuous-valued automated quality labeling algorithm using both traditional and recently developed MRS quality metrics, (2) a dual-domain (time and frequency) DL model that learns from these quality labels to assess quality scores for new data, and (3) a transient weighting algorithm informed by DL quality scores. The labeling algorithm was used to produce quality labels focused on retaining GABA peak shape in difference spectra (1) to train the DL model (2). The DL model quality scores were used to assign weights (3) for transient pairs within the final average difference spectrum. Results were compared to an existing software weighting algorithm for transient averaging and traditional transient averaging.

**Results:** Retaining only GABA-edited transient pairs with quality labels above zero, defined by metrics evaluating peak shapes, resulted in overall better traditional and recently developed mean metric values as well as better visual assessment of GABA and Glx peaks. Applying the trained DL model to *in vivo* scans, the average difference spectra calculated from the DL quality scores and weighting algorithm resulted in lower fit errors than averaging all transients with equal weights.

**Conclusion:** The proposed framework can optimize transient averaging based on quality for edited-MRS.

## 1. INTRODUCTION

In single-voxel spectroscopy, many transients are acquired to improve signal-to-noise ratio (SNR). This is particularly necessary for J-difference editing (i.e., GABA-edited MRS) in which SNR is further limited due to the subtraction of edit-On and edit-OFF difference pairs. As a result of the SNR challenges in GABA-edited MRS, 200 to 320 transients (100 to160 transient pairs) are often collected and averaged to improve SNR [1-3].

Spectral quality of the final averaged spectrum (unedited acquisitions) or average-difference spectrum, is often indicated by one or many of the following established metrics: linewidth (LW - full width at half-max), SNR, fit error, Cramer-Rao lower bound (CRLB) [4], though fit error/CRLBs are more relevant to individual metabolites rather than overall quality. Visual inspection is also often used in conjunction with quantitative metrics. While not an exhaustive list, visual inspection of quality may assess the presence of subtraction artifacts, residual water signal (including tails) resulting from poor water suppression, broadened peaks from participant motion, or the presence of other known artifacts such as spurious echoes and eddy currents [4-7]. Studies often use a combination of visual identifiers and established quantitative metrics for the purpose of keeping or discarding data from a study.

In addition to signal averaging to improve SNR, sophisticated preprocessing pipelines are now implemented to improve the quality of the final averaged spectrum. Included in the recommended preprocessing pipeline is the rejection of corrupt averages [8]. However, binary decision-making to simply retain “good” quality transients and reject “corrupt” transients is not in keeping with the notion that quality should be interpreted as a continuous range. As such, rather than simply averaging collected transients, weighing transients based on their qualitative contribution towards the final average (or difference) spectrum is a more desirable approach. Indeed, for a completely corrupt transient, a potential weighting of zero would be equivalent to transient rejection.

With the rise of machine learning in the field of MRS [9], many researchers have begun leveraging it to create quality filtering models [10-16]. While successful, many of these approaches are limited to manual subjective quality ratings [10-15], require large amounts of diverse *in vivo* data [10-15], and model quality as a binary or discrete class output (i.e., good or poor) [10-16].

We propose to develop a data-quality-based approach to weighing transient pairs in the final average GABA-edited difference spectrum. In this work, we present a three-module framework to perform transient quality evaluation that can be integrated within the preprocessing pipeline. We hypothesize that a deep learning (DL) model can be trained to assess the quality of individual transients (or transient pairs) and improve transient averaging compared to traditional methods that rely on simple averaging of all spectra.

## 2. METHODS

When acquiring GABA-edited MRS data, GABA is the primary metabolite of interest; however, the co-edited Glx (Glutamate + Glutamine) peak is often also used to evaluate overall spectral quality [17,18]. With this framework (Figure 1), we train a DL model to assess transient pair quality and weigh difference transient pairs based on their contribution to maintaining the shape of the GABA and Glx peaks in the final average GABA-edited difference spectrum. While each component of the framework is used during the initial training and validation stages, when applied to new data, only the quality assessment DL model and weighting algorithm are used together as a single pipeline. The following sections 2.1-2.3 will explain each component of the framework and its role in achieving a better-quality final average GABA-edited difference spectrum.

**Figure 1.**
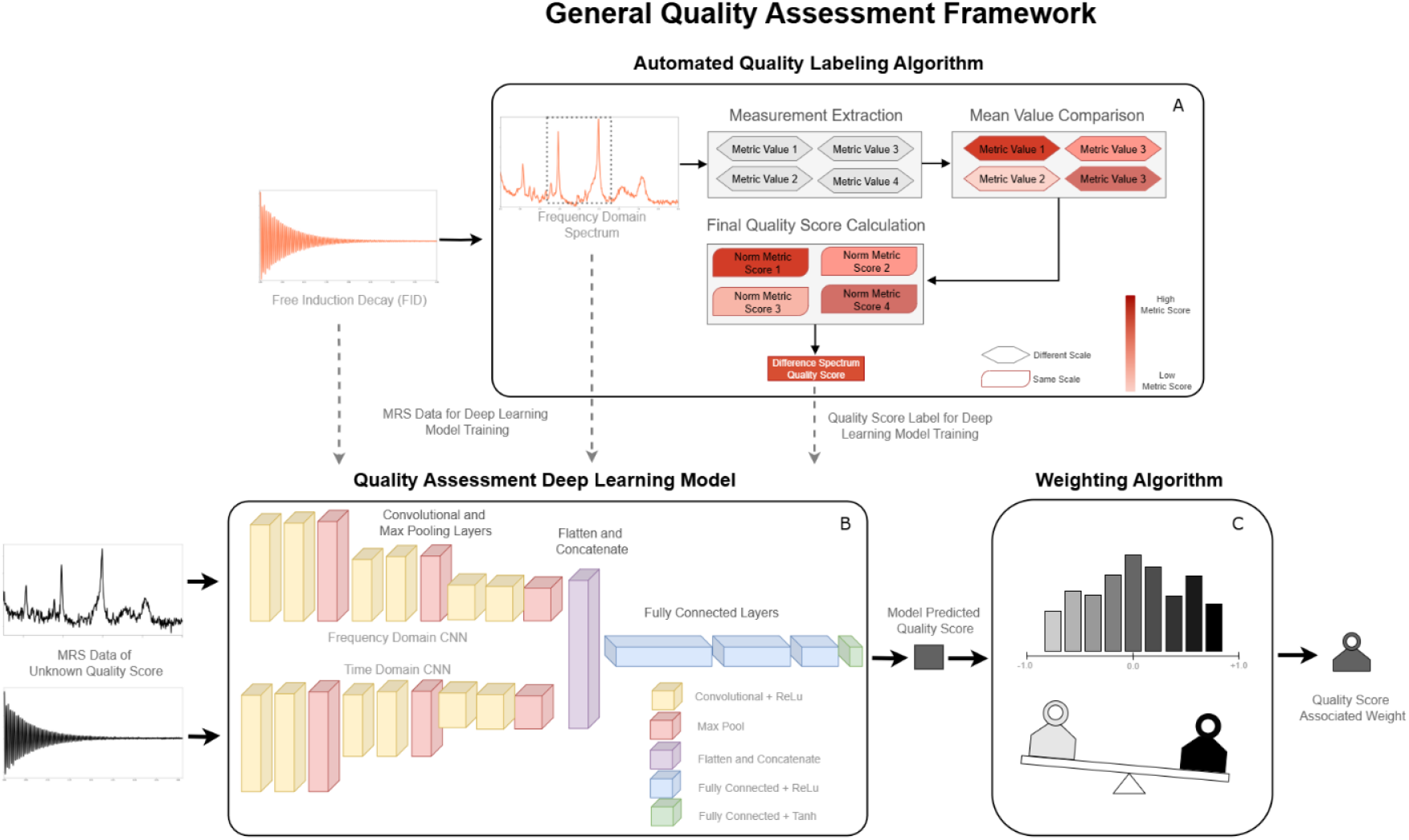
The quality assessment framework is composed of three modules. (A) Automated quality score labeling algorithm which assesses a quality score between –1 and +1 to simulated data using both traditional and/or proposed metrics. (B) Quality assessment deep learning model architecture which is trained on data from the automated labeling algorithm to predict quality scores of new unseen data. (C) Weighting algorithm which uses model predicted quality scores from the DL to assess a weight in the final average spectrum.

### 2.1 Automated Quality Labelling

#### 2.1.1 Distance Away from Mean (DAM) Labeling Algorithm

To create a supervised quality assessment model for GABA-edited MRS data, training labels of spectral quality need to be generated. Identifying good and poor-quality spectra has taken on a variety of forms, most often relying on minimum metric thresholds [19] and expert spectroscopists’ visual assessments [4]. However, these do not always account for differences in realistic achievable data quality of different participant populations (*e*.*g*. children vs. adults, clinical disorders which limit the ability to stay still), and visual inspection can vary widely based on an individual researchers’ experience or biases.

To increase consistency and reduce the subjectivity of spectral quality assessment, an automated labeling algorithm, the *Distance Away from Mean (DAM)* algorithm, was created to produce continuous quality scores between -1 (unfavorable) and +1 (favorable) which can be used as quality labels. As some of the available metrics (Table 1) require ground truths or idealized spectra, simulated data is favourable when using DAM. The DAM algorithm can label a single transient or a difference transient (the subtraction of an edit-on edit-off pair). Labeling can be based on a single quality metric or a combination of metrics, as described in Table 1. For the calculation of metrics, all data is considered in the frequency domain, i.e., after the raw FID data has been Fourier transformed.

**Table 1.** Metrics with description and example use case that can be used to calculate the DAM score.

| **Metric** | **Justification** | **Description** | **Goal** | **Example Ranges for Metabolite Targeting** |
| --- | --- | --- | --- | --- |
| Linewidth | Traditional MRS quality metric | Full width at half maximum (FWHM) | Minimize | Single Transient:<br>Creatine<br>[2.8 - 3.2 ppm]<br>Difference Transient:<br>GABA [2.8 - 3.2 ppm] |
| Signal-to-Noise Ratio (SNR) | Traditional MRS quality metric | Peak height divided by 2 times the standard deviation of noise | Maximize | Single Transient:<br>Creatine<br>[2.8 - 3.2 ppm]<br>Difference Transient:<br>GABA [2.8 - 3.2 ppm] |
| Artifact Corruption Score | Optimization of prior knowledge | Cumulative score based on number of artifacts present | Minimize | Will vary based on individual experiments (overall spectrum) |
| Peak Alignment | Translation / quantification of visual assessment | Difference between measured and ideal peak frequency<br>*ground truth required | Minimize | Single Transient:<br>Creatine<br>[2.9 - 3.1 ppm]<br>Difference Transient:<br>Inverted NAA<br>[1.8 - 2.2 ppm] |
| Subtraction Artifact Grade [20] | Translation / quantification of visual assessment | Comparison of the variance of the Choline region between difference spectra<br>*ground truth required | Minimize | Difference Transient (only):<br>Choline<br>[3.150 - 3.285 ppm] |
| Shape Score [17] | Translation / quantification of visual assessment (validated for DL in MRS) | Correlation coefficient between measured and ideal spectrum region<br>*ground truth required | Maximize | Single Transient:<br>Creatine and Choline<br>[2.8 - 3.4 ppm]<br>Difference Transient: |
|  |  |  |  | GABA and Glutamate/Glutamine (Glx) [2.8 - 3.9ppm] |
| Percent of Outlier Points | Interpretation of traditional MRS quality metric | Number of points considered greater than X standard deviations from the ideal<br>*ground truth required (can be adapted to the mean) | Minimize | Will vary based on individual experiments (overall spectrum) |
| Mean Squared Error (MSE) | Traditional Machine Learning Metric | Mean squared error of the spectra points with respect to a reference<br>*ground truth required | Minimize | Will vary based on individual experiments (overall spectrum) |

The calculation of the DAM quality scores is summarized in Figure 2. The selected metrics from Table 1 are calculated for each transient or difference pair that forms the difference spectrum. (Note, we refer to transient as the raw exported data, recognizing that some acquisitions average transients (for example across a phase cycle) prior to export off the scanner, where for this manuscript, this will also be referred to as a transient). Each individual transient metric value is either subtracted from the mean metric across all transients or difference pairs for minimization metrics (refer to Table 1) or the mean value is subtracted from the individual transient or difference pairs metric value for maximization metrics (see equation 1) to create metric scores.

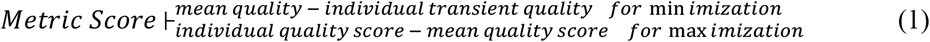

**Figure 2.**
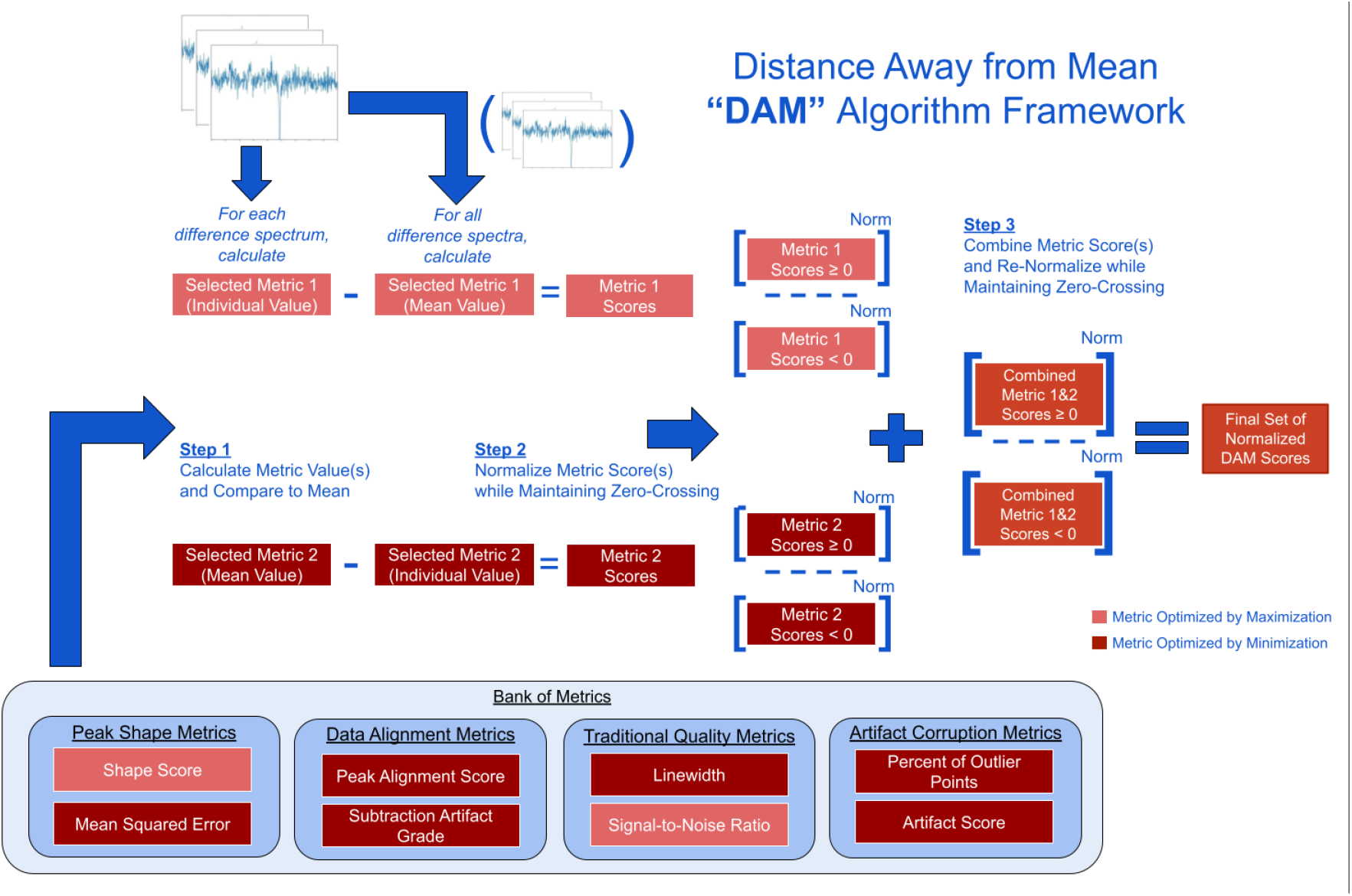
Distance Away from Mean (DAM) algorithm framework. Metrics from the bank of metrics are selected based on each project’s definition of ‘quality optimization’. Each selected metric is measured for each individual transient (or transient pair) frequency domain spectrum and compared to the mean metric value across all individual transients (or transient pairs). Positive metric scores indicate an individual transient (or transient pair) is likely to improve the average spectral quality, whereas negative metric scores indicate the individual transient (or transient pair) is likely to deteriorate the spectrum. Ranges above and below zero are normalized independently to maintain zero-crossing and a total range of [-1, +1] for each metric. Metrics are then combined (with or without weighting) and are normalized again while maintaining the zero-crossing to create the final set of normalized DAM scores.

When calculated for a single scan, this produces an intuitive result; if a metric score for a single transient or difference pair is greater than zero, it is likely of good quality and favourable to include in the final average spectrum of a scan. In contrast, if a metric score is below zero for a single transient, it is likely poorer quality and less favourable and should be downweighed in the final average spectrum. Prior to combining transient quality metric scores across multiple metrics, metric scores above 0 and below 0 are normalized independently from –1 to 0 and 0 to +1 respectively. This ensures that the magnitude of values from each metric used is comparable (if multiple metrics from Table 1 are used) and allows the algorithm to maintain the zero-crossing, separating favorable from unfavorable data. When multiple metrics are used, these normalized metric scores are summed with the opportunity to weigh each normalized metric score differently if one is to have greater importance. The combined metric scores are then normalized again across either the entire scan, in the context where DAM is only applied to a single scan, or across the entire dataset, in the context of using a dataset to train a deep learning model, using the same strategy with scores normalized independently from –1 to 0 and 0 to +1 respectively. This produces a final score between –1 (unfavorable) to +1 (favorable) for each individual transient (or transient pair).

#### 2.1.2 GABA-Edited DAM Implementation

To assess GABA-edited MRS quality, metrics measuring the accuracy of GABA and Glx peak shapes were selected. From Table 1, shape score and the percent of outlier points were selected as complimentary DAM metrics. As previously implemented [17], shape score was measured as the weighted sum of the correlation of the GABA peak between 2.8 and 3.2 ppm (weighted as 60%) and the Glx peak between 3.6 and 3.9 ppm (weighted as 40%). For this work, a secondary shape score variation, called ‘ShapeScore+L’, additionally considers the presence of lipid/macromolecule (MM) contamination between 0.8 and 1.8 ppm, which, while undesirable, often occur in voxels placed near the skull, where terms are weighted at 20% for MM, 50% for GABA and 30% for Glx. The percent of outlier points was defined as the number of spectral points which are greater than 1.5 standard deviations from the scan average divided by the total number of spectral points between 0 and 4.5 ppm. While shape score more clearly meets the objective of retaining GABA and Glx peak shape, percent of outlier points was selected as complimentary to mitigate other artifacts that may not be well captured by shape score such as baseline variations. Both metrics were set to have equal importance (each worth 50%) in the final set of DAM quality labels. A single DAM score label was assessed per difference transient.

Three DAM variations were created:

- ‘ShapeScoreOnly’: uses only the 60% GABA/40% Glx shape score metric with scores normalized per dataset.
- ‘ShapeScore+PercentOutliers’: uses the 60% GABA/40% Glx shape score and the percent of outlier points with scores normalized per dataset.
- ‘ShapeScore+L+PercentOutliers’: uses the 50% GABA/30% Glx /20% MM shape score + L variant and the percent of outlier points with scores normalized per dataset.

DAM scores were compared and normalized per set to account for varying degrees of scan quality. This prevents scores from scans with predominantly poor-quality transient pairs from being scored similarly or better than transient pairs from scans with predominantly good-quality data if normalized per scan.

To evaluate the DAM algorithm, scans were simulated with differing quality and artifacts (further described in section 2.1.2). For each DAM algorithm variation, the scan’s average difference spectrum was calculated by including only transient pairs with DAM score labels above 0 weighted evenly and compared to a Control spectrum which included all transient pairs (i.e., 160 difference pairs) weighted evenly. Average difference spectra across DAM algorithm variations and Control were visually compared to ensure DAM score labels appropriately match defined visual quality criteria. Secondly, for each DAM algorithm variation and Control, the mean and standard deviation for a combination of metrics was measured across all scans average spectra along with the percent of scans (frequency) with better metric values (as determined by whether it is a maximization or minimization metric) was also calculated.

### 2.2 DL Quality Model

#### 2.2.1 DL Quality Assessment Model Architecture and Training

As seen in Table 1, some metrics require ground truth data for their calculation. Using simulated data, DAM scores can serve as labels to train a DL model to predict quality scores for *in vivo* data, circumventing the need for ground truths *in vivo* data. To learn the association between the DAM quality score labels and spectrum quality, a DL model architecture was created (Figure 3). The model is provided with both time and frequency domain data as different artifacts can be better identified in different domains by making use of complementary information. In addition, preliminary testing showed combining information from both domains by concatenating features as opposed to a secondary model to learn the best combination of quality score predictions from each domain resulted in superior performance.

**Figure 3.**
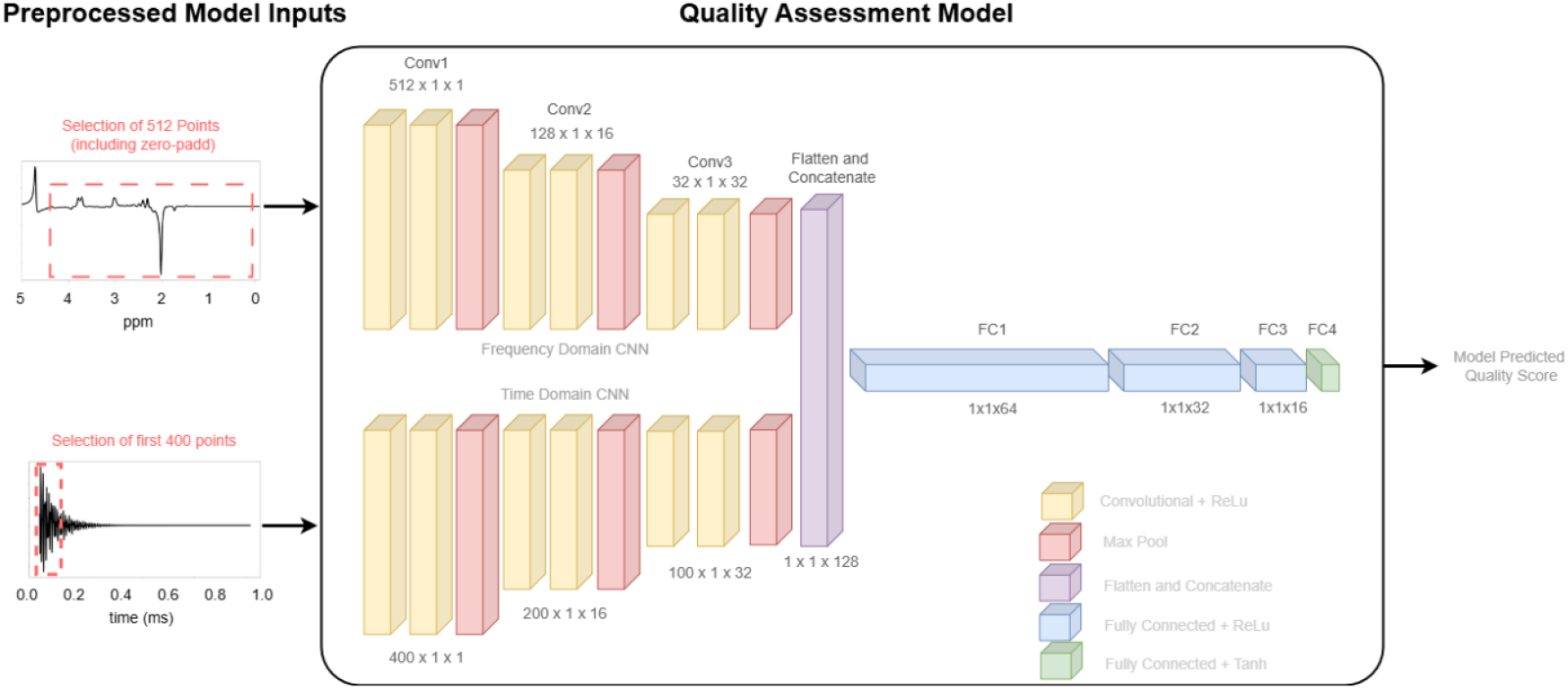
DL quality assessment model with a GABA-edited difference spectrum input example. The edited-difference spectrum is restricted to a window of 512 points and a difference FID containing the first 400 points is processed through different branches, top and bottom respectively, of the quality assessment model. Information from both domain representations is combined and used to produce a predicted model quality score.

Shown in Figure 3, the model’s top path takes a window of 512 real-valued points from a single frequency domain transient (or transient pair) converted from the time domain data using the Fourier transform. The model’s bottom path takes 400 real-valued points from the same transient (or transient pair) time domain free induction decay (FID) representation. The window of selected values is dependent on which area of the spectrum/FID the quality assessment should focus on with smaller windows of interest possible by zero-padding edges. Both representations are independently normalized prior to entering the model. Each path is processed through paired convolutional layers followed by a max pooling layer with this process repeated three times. Each path’s representation is subsequently batch normalized, flattened and concatenated into a single representation. This representation is then passed through a series of three dense layers with ReLU activation followed by a final dense layer with Tanh activation. The resulting output is a single continuous-valued quality score prediction between –1 and +1 associated with a single transient (or transient pair), i.e, equivalent to the normalized metric score label from the DAM algorithm described in section 2.1.

During training, optimal model training parameters were found to be the following: 150 epochs with early stopping based on changes to the validation error, batch size of 20, a dropout rate of 60%, Mean Absolute Error (MAE) loss function, and Adam model optimizer with a decaying learning rate of 0.001 which decreases by half every 15 epochs.

#### 2.2.2 GABA-Edited DL Model Implementation

For DL model optimization, the time domain difference FID window was selected as the default first 400 points with no zero-padding as the most important information is typically encoded within the first points of the FID. For the frequency domain difference spectrum, a spectral window between 0.00 to 4.50 ppm was selected to ensure the presence of key peaks (GABA and Glx) and best reflect the metric ranges used for the DAM algorithm score labels used for training with the remaining points zero-padded. To normalize the FIDs, each was divided by their maximum value while spectra were normalized by dividing by their 90^th^ percentile value. Similar to DAM, the DL model predicts a single quality score for each GABA difference transient pair.

The model was trained until the best validation MAE was encountered. The quality assessment model was evaluated using transient pairs from the simulated test dataset to obtain predicted quality scores. DAM algorithm score labels for the simulated test set were sorted in ascending order and plotted with the matching or paired DL model assessed quality score plotted overtop. The MAE of DL model quality scores were compared to DAM algorithm score labels per sub-range (with DAM algorithm score labels arranged from -1.0 to –0.5, -0.5 to 0.0, 0.0 to 0.5, 0.5 to 1.0). The frequency of DL model scores with signs which matched their corresponding DAM label score *(e*.*g*. positive-positive or negative-negative vs. positive-negative or negative-positive) was also calculated.

### 2.3 Transient Weighting for Final Spectrum Averaging

#### 2.3.1 Weighting Algorithm

The final module of the framework consists of a weighting algorithm, as seen in Figure 1C, to optimize the use of the DL model quality scores. The weighting algorithm dictates each individual transient’s (or transient pair’s) contribution towards the calculated final average spectrum.

The algorithm works by first sorting all individual transients (or transient pairs) by their quality score in ascending order. It then assigns the weight of the lowest quality scoring individual transient (or transient pair) to be one over the total number of transients (transient pairs) within the scan. Subsequent transients are then weighted as the weight of the previous transient (or transient pair) plus the difference in the DL model quality score. Transients (or transient pairs) with quality scores above 0, known as those above the zero-crossing, have an additional factor, known as the difference factor, multiplied against the difference in quality scores. This maximizes the difference between the weights of transients (or transient pairs) above and below the zero-crossing with a larger difference factor driving the weights of transients with quality scores below the zero-crossing towards zero. Once all weights are assigned, they are normalized to sum to one.

#### 2.3.2 GABA-Edited Weighting Algorithm Implementation

Early investigation showed that a difference factor of 100 was well-suited to maximize the difference between transient pairs with DL scores above and below 0 with difference observable in the final average GABA-edited difference spectrum.

The weighting algorithm was validated along with the framework as a whole using *in vivo* data comparing results from identical preprocessing pipelines with different weighting algorithms. Each *in vivo* transient pair for each scan was passed through our trained DL model to obtain a quality score followed by a weight from our weighting algorithm. The final average difference spectrum was then calculated using the weightings for each transient. Two additional weighting methods were compared: Gannet’s mean squared error weighting algorithm (‘MSE’) [21] and traditional weighting where all transient pairs are weighted equally (‘Control’). For fair comparison, all data was preprocessed identically and fitted in Gannet 3.3.2 [21] with only the transient weighting differing across all three methods. SNR, linewidth, fit error and GABA/Cr ratio were compared between the proposed framework and the MSE and Control methods. Appropriate statistical tests based on distribution type, parametric (t-test) or non-parametric (Wilcoxon signed rank test), were used to compare metric values from the difference spectra generated by the proposed framework compared to the MSE and Control methods with an alpha value of 0.05.

In addition, to identify which regions of the spectra were most influential to the DL model’s quality assessment, Gradient-weighted Class Activation Mapping (Grad-CAM) [22] technique was applied to the last convolutional layer in the frequency domain portion of the model.

### 2.4 Data

Traditionally, poor quality *in vivo* data containing artifacts were discarded limiting the amount of available *in vivo* data. Therefore, simulated data was used for model training and *in vivo* data for testing.

#### 2.4.1 Simulated Data

One thousand GABA+ baseline spectra (ground truths) were generated using FID-A software [23] and divided into sets of 600 for training, 200 for validation and 200 for testing. Each baseline spectra contained 22 metabolites (alanine, ascorbate, aspartate, beta-hydroxybutyrate, Cr, GABA, glutamine, glutamate, glutathione, glycine, glycerophosphocholine, glucose, myo-inositol, lactate, N-acetylaspartate, N-acetylaspartateglutamate, phosphocholine, phosphocreatine, phosphoethanolamine, scyllo-inositol, serine, taurine) [24] as well as commonly captured macromolecules (MM09, MM12, MM14, MM17, MM20) and lipid (Lip20). The ground truth spectra metabolite concentrations randomly varied by +/-10% from their averaged reported concentrations in literature. The acquisition parameters of the simulated datasets were: magnetic field strength (3T), MEGA-PRESS variant (FID-A’s MegaPressShaped_fast [23,25]), 14 ms editing pulses at 1.9 ppm and 7.46 ppm, alternating editing pulse location at each TR, TR/TE = 2 s/68 ms, spectral width (5000 Hz), and the number of spectral data points (N = 4096).

Each baseline spectrum was used to generate a set of 320 transients (160 ON and 160 OFF). With the DAM algorithm normalizing scores in the final stage based on all difference spectra in the entire dataset, maximizing the diversity of quality defined by none, one or multiple artifacts is important. The *SMART_MRS* toolbox [26] was used to add different combinations of artifacts and noise with varying parameters based on what has been commonly observed in GABA-edited MRS *in vivo* data. Artifacts were added to approximately half of the transients from each scan which included complex noise with a standard deviation between 1 and 3.5×10^−5^, frequency and phase shifts uniformly distributed as ±15 Hz and ±180 degrees, lipid/MM peaks appearing between 0.8 and 1 ppm, spurious echoes, eddy currents, line broadening, and baseline distortions.

All simulated transients generated were passed through the DAM algorithm. Both the training and validation sets were balanced according to six sub-ranges of score values (between -1.0 to – 0.5, -0.5 to –0.25, -0.25 to 0, 0 to 0.25, 0.25 to 0.5, 0.5 to 1.0). Based on the ideal number of samples per sub-range, the greatest common factor for the number of scores within each range was found and used to preliminary fill our balanced dataset. All remaining samples were subsequently collected from ranges with data available to create the most balanced dataset.

#### 2.4.2 In Vivo Data

34 MEGA-PRESS data sets composed of 64 difference transients [27] were used for the weighting algorithm validation (section 2.3.2). The acquisition parameters were: magnetic field strength (3T), 14 ms editing pulses at 1.9 ppm and 7.46 ppm, alternating editing pulse location at each TR, TR/TE = 1.8 s/68 ms, spectral width of 5000 Hz, and 4096 spectral data points. In addition, a single 320 transient scan from a publicly available dataset [28] were used with acquisition parameters: magnetic field strength (3T), MEGA-PRESS, 15 ms editing pulses at 1.9 ppm and 7.46 ppm, alternating editing pulse location at each TR, TR/TE = 2 s/68 ms, spectral width of 5000 Hz, and 4096 spectral data points. The above was selected for its varying quality and artifacts. Scans were retained (as opposed to shuffling transients) to enable the measurement and comparison of quality metrics for entire scans.

## 3. RESULTS

### 3.1 GABA-Edited DAM Algorithm Evaluation

Figure 4 shows three sample average spectra which include only transient pairs with DAM quality score labels above 0 from each of the three DAM variations. Additionally, Control, which includes all transients, is shown as well as the Ground Truth simulated spectrum (i.e., prior to noise or artifacts being added). Control and ShapeScoreOnly do not retain the ground truth GABA and Glx (3.5 ppm) peaks as well as ShapeScore+PercentOutliers and ShapeScore+L+PercentOutliers. The two DAM variants with multiple metrics, ShapeScore+PercentOutliers and ShapeScore+L+PercentOutliers, visually appear to outperform the DAM algorithm variant using a single metric, ShapeScoreOnly, suggesting that using multiple complimentary metrics can be superior to using a single metric.

**Fig 4.**
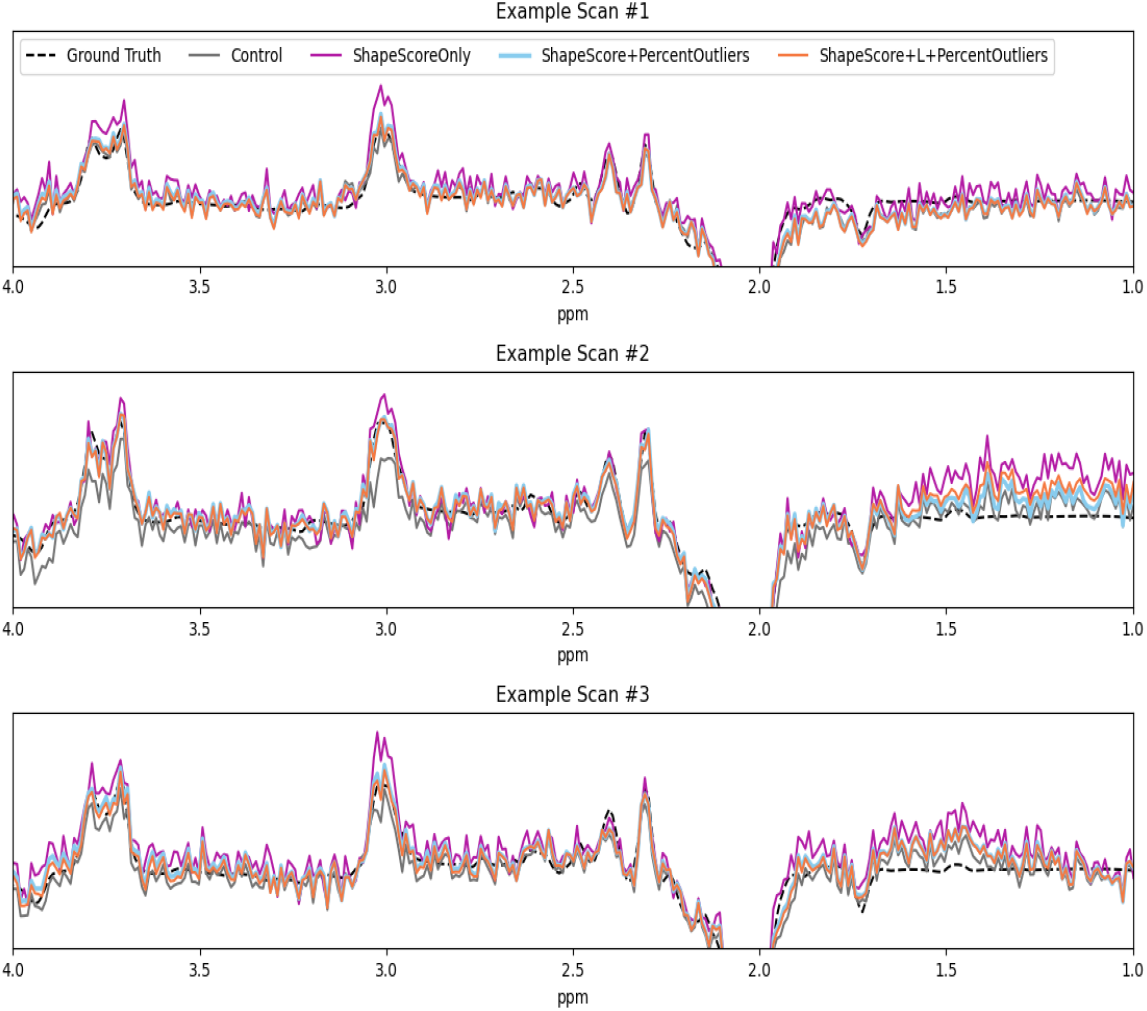
Comparison of average spectra for three scans composed only of transient pairs with DAM quality labels above zero for three DAM automated quality label algorithm variations: ShapeScoreOnly (purple), ShapeScore+PercentOutliers (blue), ShapeScore+L+PercentOutliers (orange). For comparison, the average difference spectrum with all transients is shown in grey (Control) and the simulated ideal spectrum (ground truth, prior to addition of noise and artifacts) is the black hashed line.

Table 2 compares the metric values between each DAM variation and Control for shape score, SNR, linewidth, and MSE

**Table 2.** Mean metric values of three DAM algorithm variations for simulated scans which include transient pairs with DAM quality scores above zero compared to the inclusion of all transient pairs to calculate the scan’s average spectrum (Control). Percent of scans with better metric values compared to Control are also presented. Results that are bold indicate better performance compared to Control.

|  |  | Shape Score | SNR | Linewidth | MSE |
| --- | --- | --- | --- | --- | --- |
| **Control** | Mean ± SD | 0.847 ± 0.0893 | 4.34 ± 1.06 | 0.121 ± 0.0571 | 0.0380 ± 0.0450 |
| **'ShapeScore Only'** | Mean ± SD | **0.886 ± 0.0637** | **5.78 ± 0.935** | **0.0895 ± 0.0243** | 0.128 ± 0.0826 |
|  | Percent of Scans Better than Control | **67.8%** | **94.6%** | **60.1%** | 12.7% |
| **'ShapeScore +Percent Outlier'** | Mean ± SD | **0.876 ± 0.0733** | **4.912 ± 0.714** | **0.101 ± 0.0360** | **0.0338 ± 0.0396** |
|  | Percent of Scans Better than Control | **74.1%** | **76.1%** | 45.1% | **51.9%** |
| **'ShapeScore +L +Percent Outlier'** | Mean ± SD | **0.874 ± 0.0747** | **4.80 ± 0.692** | **0.0966 ± 0.0352** | **0.0337 ± 0.0393** |
|  | Percent of Scans Better than Control | **71.9%** | **71.4%** | 45.2% | **52.2%** |

The ShapeScoreOnly DAM variant shows both the greatest improvements in SNR, linewidth and shape score while also showing the greatest decrease in mean squared error. This is supported by Figure 4 which shows that while the ShapeScoreOnly variation produces spectra which moderately retains peak shapes, it appears to systematically overestimate peak amplitude compared to other model spectra. When comparing the ShapeScore+PercentOutliers and ShapeScore+L+PercentOutliers variants, both perform similarly on all metrics, with only linewidth showing a small drop in performance compared to Control. Given the potentially wider applicability of the ShapeScore+L+PercentOutliers variant to the presence of MMs/Lipids, this was selected as the best DAM algorithm variation.

### 3.2 GABA-Edited DL Quality Model Evaluation

To evaluate the ability of the DL quality model to learn labels from the ShapeScore+L+PercentOutliers DAM algorithm variation, Figure 5 shows the DAM label scores sorted and plotted in ascending order with its paired (matched) DL model quality score plotted over top for each difference spectrum in the simulated test set. Subregions of DAM quality label scores (−1.0 to -0.5, -0.5 to 0.0, 0.0 to 0.5, and 0.5 to 1.0) are indicated with the corresponding MAE and the rate of correct sign. The overall MAE of the test set was 0.123 with the highest MAE in the 0.0 to 0.5 subregion with a MAE of 0.144. The rate of correct sign (positive or negative) shows how often the model predicted quality score matched the sign of the DAM score label, with the lowest value being 78.35% in the -0.5 to 0.0 subregion.

**Figure 5.**
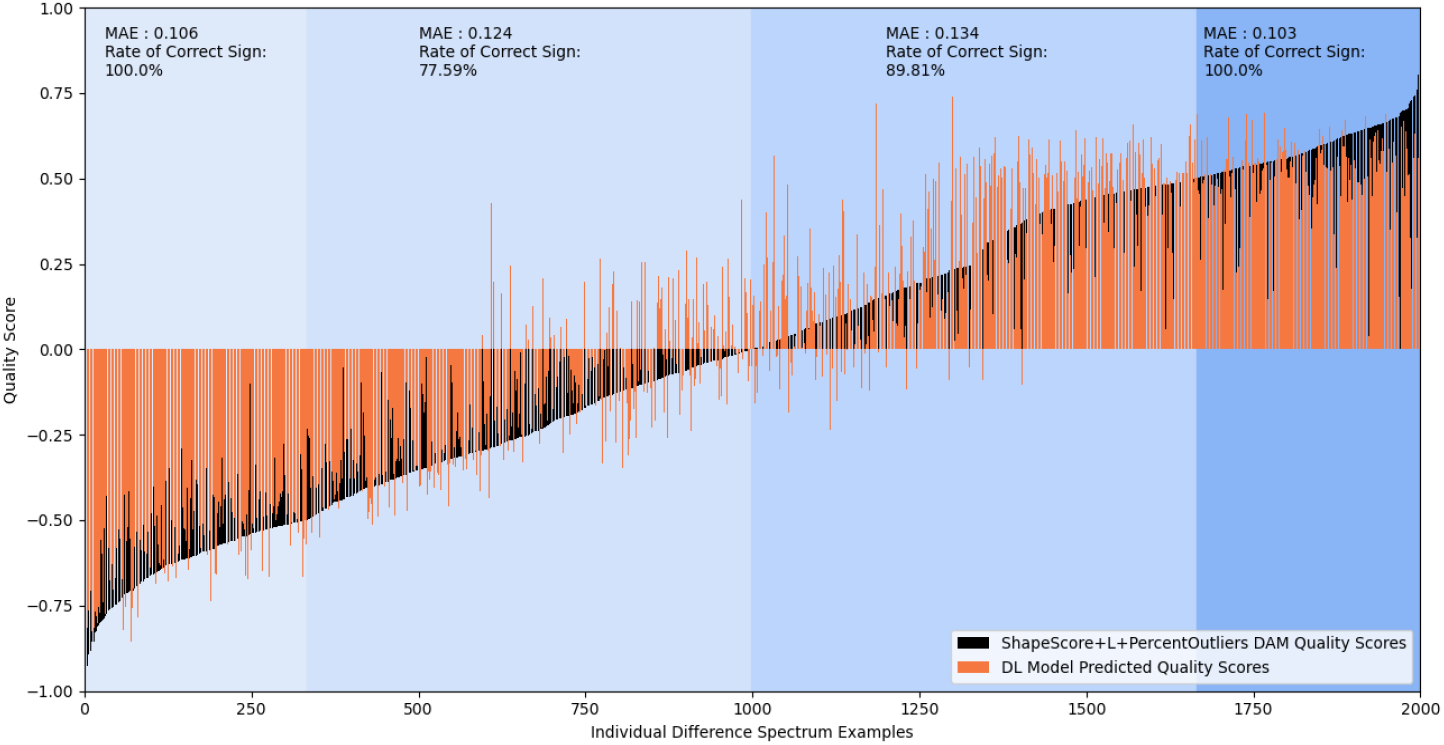
Evaluation of the quality assessment DL model predicted s cores (orange) compared to the true DAM (ShapeScore+L+PercentOutliers) score labels (black) with simulated samples ordered from lowest to highest true DAM label score. Scores are divided into 4 sub-ranges (−1:-0.5, -0.5:0, 0:0.5, 0.5:1) with magnitude of the MAE and the rate of correctly predicted score sign (positive/negative) measured for each.

### 3.3 GABA-Edited Framework Evaluation

Subsequent to simulated data validation of the automated labeling algorithm and DL model, trained using ShapeScore+L+PercentOutliers DAM algorithm variation, the framework was evaluated on *in vivo* data. Figure 6 shows three examples of in vivo spectra comparing our framework, consisting of the DL model quality scores implemented as transient pair weights using our weighting algorithm, compared to software MSE weighting and Control’s equal transient weighting. Our framework appears to downweigh transient pairs that will contribute to subtraction artifacts while up weigh transient pairs that will contribute to better SNR.

**Figure 6.**
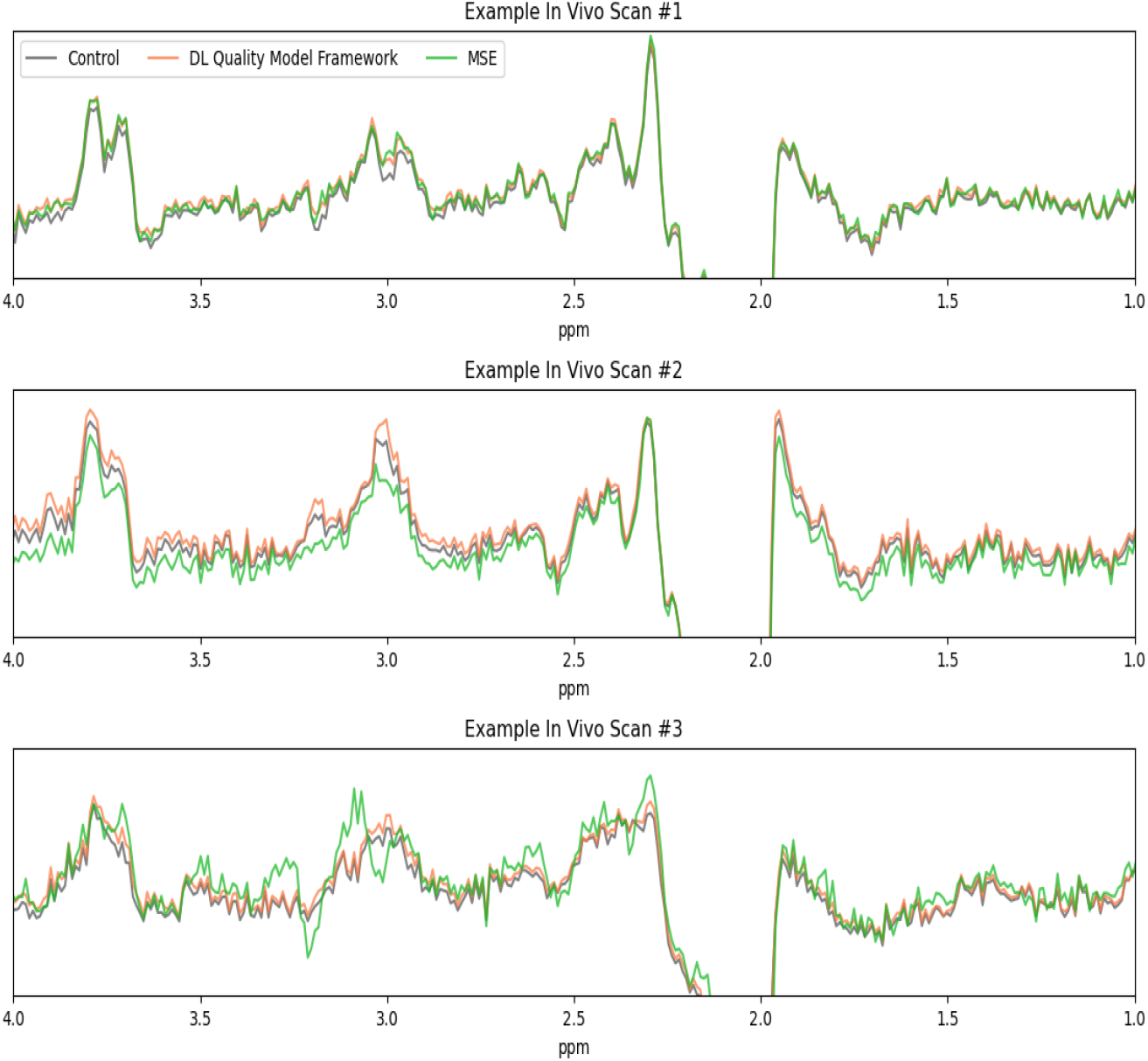
Evaluation of our quality framework with *in vivo* sample scans from three averaging methods where transient pairs were weighted according to our DL quality model framework (orange), all transient pairs were weighed equally (Control (grey)), or all transients were weighted according to leading software (MSE (green)).

In Figure 7, SNR, linewidth, fit error, and GABA/Cr ratio for all *in vivo* average spectra were compared across three averaging methods: 1.) Our DL quality model framework (the DL model assigned quality scores converted to weights for each transient pair to calculate the average spectrum), 2.) Control (equal weights across all transients to calculate the average spectrum), and 3.) MSE (weights based on MSE, default in Gannet [21] to calculate the average spectrum). Data normality testing showed that SNR, fit error and concentration had non-normal distributions and therefore a Wilcoxon signed rank test was used to assess significant differences between the performance of our framework compared to ‘Control’ and ‘MSE’, while linewidth had a normal distribution and therefore used a paired t-test to assess significance.

**Figure 7.**
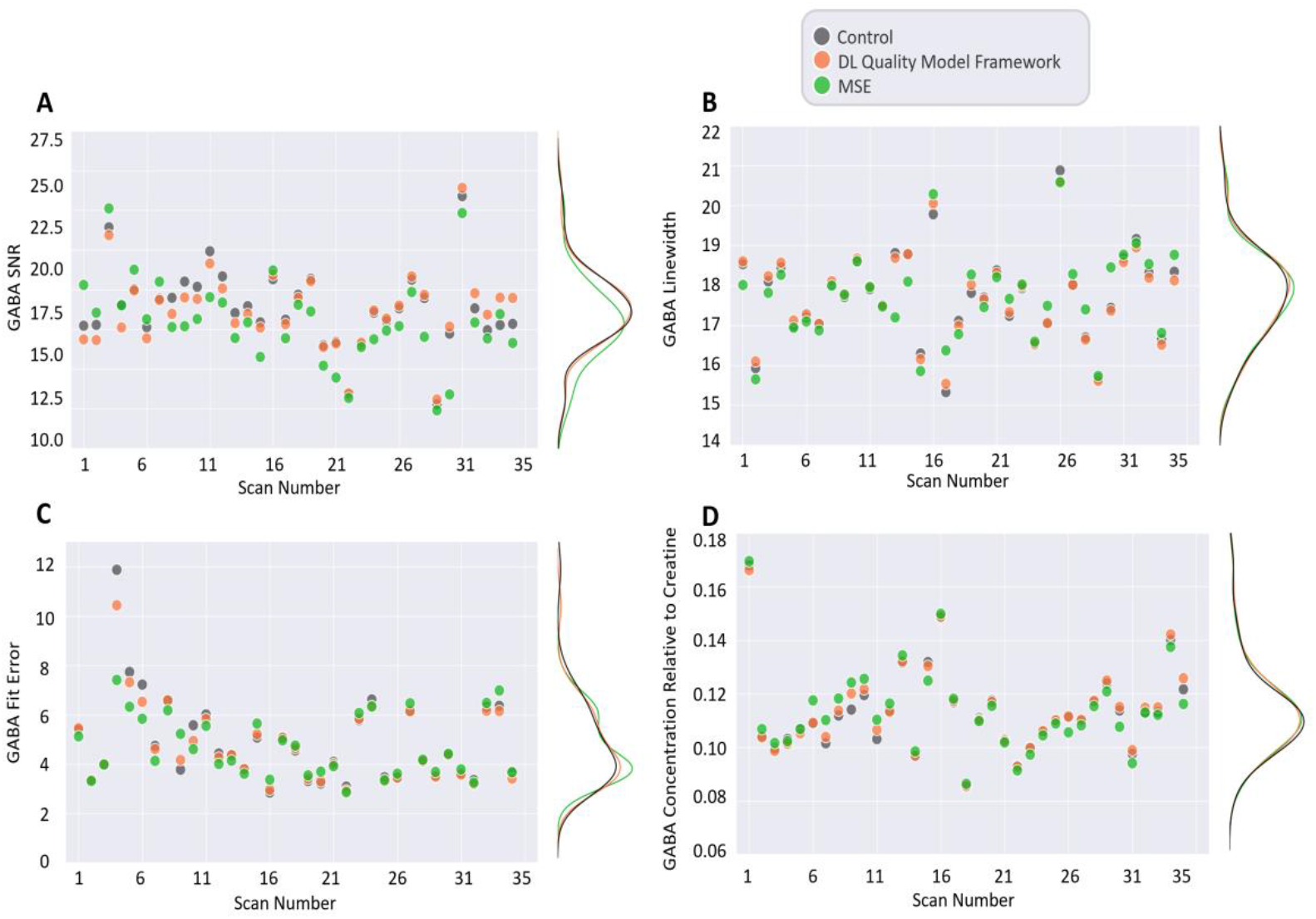
Evaluation of our DL quality model framework (orange) on *in vivo* data compared to ‘Control’ (equally weighted transients, grey) and ‘MSE’ (MSE assessed weights via software, green). Scatter plots of the test set scan number by (A) GABA SNR, (B) GABA linewidth (FWHM), (C) GABA fit error, and (D) GABA concentration relative to Creatine with probability density distributions to the right of each subplot.

SNR was significantly higher when using the DL quality model framework compared to MSE (p = 0.007) with no significant differences compared to Control (p = 0.334). Fit error and GABA/Cr ratio were not significant between the DL quality model framework and MSE (p = 0.994 and p = 0.852). However, the DL quality model framework showed a significantly smaller fit error (p = 0.015) and larger GABA/Cr (p = 0.027) than Control. There were no significant differences in GABA linewidth.

While typically only single transient pairs are passed through the DL model, for visualization purposes, Figure 8 shows the saliency maps for three final average difference spectra examples. Regions of the spectrum indicated by warmer colors, such as the GABA, Glx and MM/Lipid peak, indicate areas of high importance within the DL model when determining quality.

**Figure 8.**
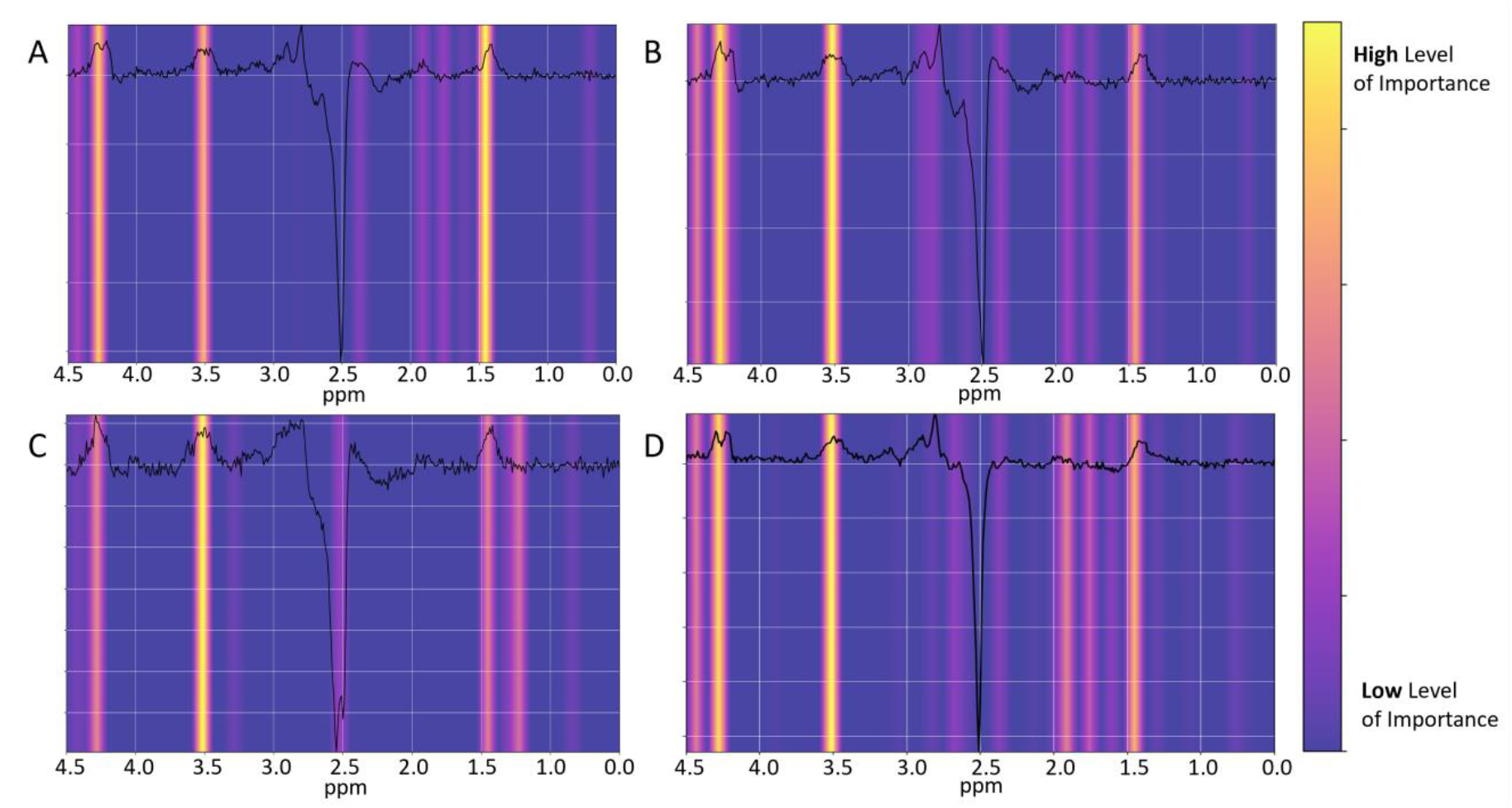
Saliency maps overlayed on four example *in vivo* final average difference spectra passed through our quality assessment DL model trained on our DL quality model. Areas of high importance for model determination of quality are indicated by warm colors while areas with low levels of importance are indicated by colder colors.

## 4. DISCUSSION

The objective of this work was to improve GABA-edited MRS data quality through the optimization of transient pair averaging within the final averaged spectrum. To accomplish this, we developed a three-module framework consisting of 1) an automated quality score labeling algorithm (DAM), 2) a DL model for transient quality scoring, and 3) a weighting algorithm to translate the DL model quality scores to transient pair weights within the final average difference spectrum. The code is publicly available for reproducibility purposes.

The DAM algorithm metrics were selected to best retain GABA and Glx peak shapes. As shown in Figure 4 and Table 2, while using one metric, ShapeScoreOnly, (compared to multiple metrics, ShapeScore+PercentOutliers and ShapeScore+L+PercentOutliers) resulted in better all-around mean metric values, it significantly underperformed in terms of MSE which can also be seen visually with spectra which consistently overestimated peak amplitude resulting in poor retention of original peak shape, the original objective set out for the DAM algorithm, compared to DAM variants which used two metrics. Although this indicates that using more than one complimentary metric can improve DAM algorithm performance, preliminary analyses indicated that using too many metrics resulted in difficulties optimizing all metrics simultaneously.

The DAM automated labeling algorithm provides continuous, consistent, objective, and quantifiable quality metrics. However, its accuracy in scaling transient quality increases with an increasing number of samples (transients or transient pairs) and thus requires many scans to form a large dataset for optimal performance. As such, a trained DL model is a suitable alternative to provide quality scores to new transients without the need for other transients (or transient pairs) to assess quality comparatively. Simulated data testing of the DL model showed good performance with some cases indicating an overestimation of quality scores though this was determined to be acceptable as predictions still show a gradual increase, as desired, compared to true values from DAM score labels. Confidence in the DL model’s performance was also supported by Figure 8, which shows that the DL model relies most on the GABA and Glx peaks for quality determination. The 0.9 MM/lipid peak was also marked as highly important for the DL quality model predictions. This generates additional confidence in the reliability of DL-quality labeling for MRS transients. When applied to *in vivo* data, the framework (DAM training labels, DL model, and weighting algorithm) outperformed existing software transient weighting algorithms with superior SNR and traditional transient averaging with superior fit error.

In this study, we examined GABA-edited MRS data for both the DAM quality rating as well as the DL quality labeling and weighting. With minor changes to the different modules, these modules can be adapted to different data types (i.e., unedited short-echo time data). In addition (or alternatively) different quality metrics (i.e., maximize SNR or minimize linewidth compared to maximize ShapeScore) would be selected. While the framework performs well for the above GABA-edited data, we have not tested different data options as this is outside the scope of this work (with the opportunity to explore this further offered to others through our open-access code). Additionally, there is also the option of integrating preprocessing steps, such as frequency and phase correction, within the framework to better tailor the DL model’s performance. We also acknowledge the other limitations of this work. First, training on simulated data offers the possibility of using metrics which compare to a ground truth to better objectively determine improvement where the use of *in vivo* data would be limited to using the full scan average for comparison. However, *in vivo* data may better capture true artifact variance as we are only beginning to understand and note key artifact parameters. Additional training with *in vivo* data may further improve DL model performance. In addition, our GABA-edited MRS example took the approach of scoring edit-On/edit-OFF transient pairs (difference transient) as opposed to all individual transients. The approach here reduces the complexity of the DAM equation and the inputs to the DL model as the focus is on a single spectrum appearance (as opposed to transients with two conditions: edit-ON and edit-OFF). Alternatively, scoring individual transients may provide additional benefit and flexibility as individual transients are weighted independently of the quality of the transient pair.

## 5. CONCLUSION

A three-module framework for assessing edited-MRS transient quality consisting of an automated continuous quality score labeling algorithm, a flexible dual-domain DL model, and a weighting algorithm was presented. The use of the framework for optimizing GABA and Glx peak shape showed improved performance over traditional averaging and existing software weighting algorithms for averaging on *in vivo* data.

## DATA AND CODE AVAILABILITY

The framework’s code, consisting of a Python notebook that walks through each module of the framework to retrain on new data with new optimization criteria, is available through the following GitHub repository (Git commit SHA: e492ecb7a15be8a2be437860f69f0d0ff01c2e14):

https://github.com/HarrisBrainLab/GABA-Edited-MRS-Quality-Assessment-Framework

## ACKNOWLEDGEMENTS

This work was presented in part by Bugler et al. as “*Quality assessment tool using deep learning for GABA-edited MRS data*” at the International Society for Magnetic Resonance in Medicine Annual Meeting 2024 in Singapore.

